# 16p11.2 Copy Number Variation Alters Genome Architecture and Transcriptional Regulation During Neurodevelopment

**DOI:** 10.64898/2026.05.07.723480

**Authors:** Marco Bertin, Marcel Misak, Mohit Navandar, Azza Soliman, Ruxandra-Andreea Lambuta, Roshan Tumdam, Jennifer Winter, Matthias Linke, Stephan Käseberg, Malin Dewenter, Katalin Komlosi, Lea Zografidou, Oliver Tüscher, Sergiy Davydenko, Daniel Pieh, Jannik Foerster, Susanne Gerber, Susann Schweiger

**Affiliations:** Institute of Human Genetics, University Medical Center Mainz, Mainz; Department of Psychiatry and Psychotherapy, University Medical Center Mainz, Mainz; Institute for Quantitative and Computational Biosciences (IQCB), Mainz; Institute of Molecular Biology, Mainz; Leibniz Institute for Resilience Research, Mainz; Department of Psychiatry, Frankfurt; Department of Human Genetics, Bioscientia, Ingelheim; Center for Rare Diseases, University Medical Center Freiburg, Freiburg; Department of Psychiatry, Psychotherapy and Psychosomatic Medicine University Medicine Halle (Saale) of the Martin Luther University Halle-Wittenberg (MLU) & German Center for Mental Health (DZPG), partner site Halle-Jena-Magdeburg

**Keywords:** 16p11.2 CNVs, neurodevelopmental disorders, 16p11.2 microduplications, iPSCs, neurons, MAPK3/ERK pathway, 3D chromatin structure, TAD domains

## Abstract

Microdeletions and microduplications in 16p11.2 are responsible for a spectrum of neurodevelopmental disorders (NDDs) with partially reciprocal and overlapping symptoms. However, the clinical variability in 16p11.2 microduplication patients is significantly greater than that in patients carrying a microdeletion. Here, we use iPSCs derived from members of a family carrying a 16p11.2 microduplication and model neurodevelopment through *in vitro* differentiation into neural progenitor cells (NPCs) and neurons. The analysis is complemented by reanalysis of publicly available data from 16p11.2 microdeletion patients. Transcriptome analysis revealed MAPK3-centered hubs of upregulated genes in the microduplication-carrying and downregulated in the microdeletion-carrying cells, indicating that MAPK3 is a central driver of 16p11.2 Copy Number Variation (CNV) pathology. While genes within the 16p11.2 region showed about a two-fold reduction in expression in cells carrying the microdeletion, their expression levels in microduplication-derived NPCs and neurons, but not in iPSCs, were elevated to a degree much higher than expected. This observation was accompanied by a substantial number of dysregulated genes unrelated to the genes in the critical region or their interaction networks. To further investigate whether altered chromatin organization may accompany these transcriptional changes, we generated Hi-C data from patient and control iPSCs and NPCs. This revealed increased chromatin contacts within the duplicated 16p11.2 region in patient-derived iPSCs, while genome-wide compartment analysis showed that increased compartments preferentially co-occurred with upregulated differentially expressed genes, particularly in NPCs. Together, these findings suggest that 16p11.2 microduplication may influence gene expression through both local dosage-dependent mechanisms and broader, differentiation-associated changes in chromatin organization. Our data support a model in which CNV-associated genome architecture changes may modulate transcriptional dysregulation and contribute to the variable neurodevelopmental phenotypes associated with 16p11.2 rearrangements.

## Introduction

Neurodevelopmental disorders (NDDs) arise from a complex interplay of genetic and environmental factors, with genetic alterations contributing substantially to disease risk. Among these genetic factors, copy number variations (CNVs) within the human 16p11.2 genomic locus are strongly associated with a range of pleiotropic phenotypic manifestations, including autism spectrum disorder (ASD), intellectual learning disability (ID), communication disorders, psychiatric features, including bipolar disorder, anxiety disorder and schizophrenia, and altered physical appearance (Rein and Yan 2020).

The 16p11.2 gene locus is flanked by 147-kb low-copy repeats (LCRs) that are 99% identical (Kumar et al. 2008), which makes the region vulnerable to misalignments through nonallelic homologous recombination, resulting in deletion (OMIM# 611913) or duplication (OMIM# 614671) events. These CNVs span a variable length of 500-600 kb and involve 27-29 protein-coding genes. Recent literature re-evaluations indicate that the incidence of the 16p11.2 microdeletion is estimated to be approximately 0.032% in the general population (Auwerx et al. 2024a). In comparison, the rate of 16p11.2 microduplication is estimated to be approximately 0.036% (Auwerx et al. 2024a). An increased incidence is observed in NDD patients, with microdeletions found in 0.276% and microduplications in 0.254% of affected NDD patients (Auwerx et al. 2024a). Interestingly, the phenotypic effects associated with these CNVs in some domains show apparent reciprocal effects. Thus, microdeletions of the 16p11.2 locus are correlated with macrocephaly and obesity, whereas microduplications are associated with microcephaly and a lower body mass index (BMI) (Jacquemont et al. 2011; Qureshi et al. 2014; D’Angelo et al. 2016; Auwerx et al. 2024a).

While no significant differences in the prevalence of ASD, ID, or epilepsy are noted between 16p11.2 microdeletion- and microduplication-carrying patients, microduplications generally exhibit greater variability in disease phenotypes. Notably, developmental delay, language impairment, and psychiatric conditions such as schizophrenia are more common in microduplication patients than in microdeletion patients, with schizophrenia and bipolar disorders being consistently reported in microduplication patients only (Chang et al. 2017; Zhou et al. 2018; Jutla et al. 2020; Auwerx et al. 2024a).

Many genes within the 16p11.2 region play pivotal roles in diverse cellular and developmental processes. Particularly noteworthy are the *MAPK3* and *TAOK2* genes, which have been highlighted in human and mouse studies for their influential role in neurodevelopmental signaling pathways (Deshpande et al. 2017; Richter et al. 2019). Additionally, deletion studies have linked the *KCTD13* gene to neurodevelopment and shown that it regulates synaptic connectivity and influences dendritic spine morphology (Golzio et al. 2012), whereas *PRRT2* mutations in humans have been associated with seizure and movement disorders (Chen et al. 2011; Heron and Dibbens 2013). Recent studies using human-induced pluripotent stem cells (iPSCs) derived from peripheral blood mononuclear cells of 16p11.2 deletion carriers revealed an association between *MAPK3* haploinsufficiency and disrupted global transcriptomic profiles, impacting neural development at critical stages (Liu et al. 2023). In line with this, investigations in iPSC-derived neuronal cells of schizophrenia patients carrying 16p11.2 duplication have revealed dysregulated synaptic gene expression, altered calcium signaling, and impaired dendritic morphology, suggesting links to schizophrenia pathophysiology through glutamatergic dysfunction and altered neuroarchitecture (Parnell et al. 2023; Leone et al. 2024). In a study with iPSC-derived neural precursor cells and neurons of two families with a 16p11.2 microdeletion, the authors identified a disruption of the MAPK3/ERK pathway as the likely major driver for the 16p11.2 microdeletion phenotype (Liu et al. 2023).

CNVs change the size of the chromatin in a densely packed nucleus, which suggests an effect on the chromatin 3D structure and associated functions. Approximately 2 mt of DNA is stored in each nucleus with a diameter of just 5-6 µm. This is only achievable through highly compact and dynamic 3D folding of the genome (Cremer and Cremer 2010, Lieberman-Aiden et al. 2009). However, the spatial and hierarchical structure of chromatin not only allows for maximum compaction and condensation of DNA but also has a vital role in the transcriptional control of genes and in establishing direct, long-range interactions between chromosomal sequences. Spatial proximity, for example, is necessary for enhancers to regulate the transcription of target genes, and chromatin accumulation near the nuclear lamina has been linked to gene silencing and replication timing (Deng et al. 2014). In addition to long-known chromatin structures, such as nucleosomes and their arrangement, other highly organized features of genomic architecture have been reported, such as relatively small DNA loops (hundreds of kb long), larger 0.5-1.0 Mb topologically associated domains (TADs and sub-TADs), but also multi-contact chromatin hubs, cliques, and so-called “frequently interacting regions” (FIREs) (Dixon et al. 2012; Nora et al. 2012; Rao et al. 2014). These structures are stabilized by structural proteins that bind to chromatin, such as Cohesin and CCCTC-binding factor (CTCF), and are key for the fine-tuned regulation of gene expression through promoter□enhancer interactions (Wendt et al. 2008).

During nervous system development, NPCs differentiate into diverse neuronal and glial cell types. This differentiation is accompanied by extensive changes in gene expression, driven by epigenetic modifications and by dynamic reorganization of the 3D genome, which modulates enhancer-promoter interactions to fine-tune transcriptional programs. (Yao et al. 2016; Kishi and Gotoh 2018; Yoon et al. 2018; Lu et al. 2020). On the other hand, aberrant changes in chromatin architecture have also been identified as triggers for various disorders, including NDDs such as fragile X syndrome and schizophrenia (SCZ) (Sun et al. 2018; Girdhar et al. 2022). Large-effect CNVs, including microdeletions and microduplications, are thought to substantially affect chromatin structure and 3D organization, thereby significantly influencing gene expression (Rovina et al. 2020; Chilinski et al. 2022; Franke et al. 2022).

Here, we describe a family with a 16p11.2 microduplication over two generations that is associated with highly heterogeneous phenotypes in different family members. We generated iPSCs, NPCs, and neurons from the index patient and her unaffected mother and performed RNA sequencing to compare the transcriptomes. We observed progressive transcriptional alterations throughout neurodevelopment, with a marked increase in differential gene expression during neuronal differentiation. Our data show that MAPK3 is a potent regulator of these gene expression changes. We reanalyzed publicly available iPSC-derived neuronal transcriptomic data from 16p11.2 microdeletion patients (Liu et al. 2023) and identified an opposing MAPK3-centered regulatory network influencing neuronal differentiation. To investigate whether the 16p11.2 microduplication is also associated with changes in higher-order chromatin organization, we generated Hi-C data from patient and control iPSCs and NPCs. This analysis revealed increased chromatin contacts within the duplicated 16p11.2 region in patient-derived iPSCs, suggesting that the duplication may alter local chromatin organization in a cell-state-dependent manner. Genome-wide compartment analysis further indicated that increased chromatin compartments preferentially co-occurred with upregulated differentially expressed genes, particularly in NPCs. These observations suggest that transcriptional changes in 16p11.2 microduplication may reflect not only dosage-dependent effects of genes within the duplicated interval but also broader changes in chromatin organization during neural differentiation. Our study provides valuable insights into the molecular mechanisms underlying neurodevelopmental features in patients carrying 16p11.2 CNVs, illuminating the complex interplay between chromatin, genes, and biological processes crucial for brain development and function.

## Results

### Family with 16p11.2 Microduplication-Associated Psychiatric Phenotypes and Neurodevelopmental Delays

In this study, we included a German family with three members carrying a heterozygous 554 kb copy number gain on 16p11.2 (chr16: 29,622,757-30,177,240; hg19) identified via chromosome microarray analysis (CMA). Individuals exhibit a spectrum of psychiatric phenotypes and neurodevelopmental symptoms. The microduplication is found in the father, son, and daughter (Figure 1A). The duplication spans 28 protein-coding genes and five lncRNA genes, extending from *SLC7A5P1* to *MAPK3*, both of which are fully duplicated (Figure 1C). The daughter approached the genetic counseling service for the first time at the age of 29. She was diagnosed with speech development delay and moderate learning disability in childhood. During her adolescent years, she endured bullying and developed episodes of excessive anxiety and depression. At the age of 18, she experienced a psychotic episode and underwent treatment involving antipsychotic medication combined with psychotherapy. The clinical assessment revealed cognitive weaknesses, resulting in an IQ score of 62.5. She had two lumbar disc herniations without explanatory physical strain at the ages of 27 and 28 years. No further signs of connective tissue disorders were observed. Physical examination revealed no dysmorphic features, a normal head circumference (head circumference 55 cm, 15th percentile), and a normal BMI (body size 170 cm, weight 84 kg). She currently requires assistance in her daily activities and works as a volunteer once a week. The family’s son had delayed speech development and encountered learning challenges during childhood. As he transitioned into adulthood, he received a diagnosis of severe depression and a personality disturbance when he turned 18, necessitating treatment with antipsychotic medication. At the age of 21, the patient experienced acute aortic dissection. Physical examination revealed no dysmorphic features, a normal head circumference (head circumference 58 cm, 75th percentile), and a normal BMI (body size 191 cm, weight 88 kg). Together with overall joint laxity, Marfan syndrome was suspected but could not be confirmed molecularly. Later in life, at the age of 34, he was diagnosed with Hodgkin’s disease. The father has been receiving psychological treatment for depression and personality disorders since the age of 19. He had experienced sexual abuse during adolescence. He also stated that he had several herniated discs at approximately 40-50 years of age with no further signs of connective tissue disorder. Despite his mental health challenges, he was able to raise his two children together with his wife and maintain a career as a craftsman. Physical examination revealed no body abnormalities or pathological head circumference (head circumference 57 cm, 50th percentile), and the BMI was normal (body size 182 cm, weight 80 kg). Overall, psychiatric illnesses seemed to have a smaller impact on him than on his two children. Both the daughter and the mother consented to provide a skin biopsy for further molecular analysis.

**Figure 1:**
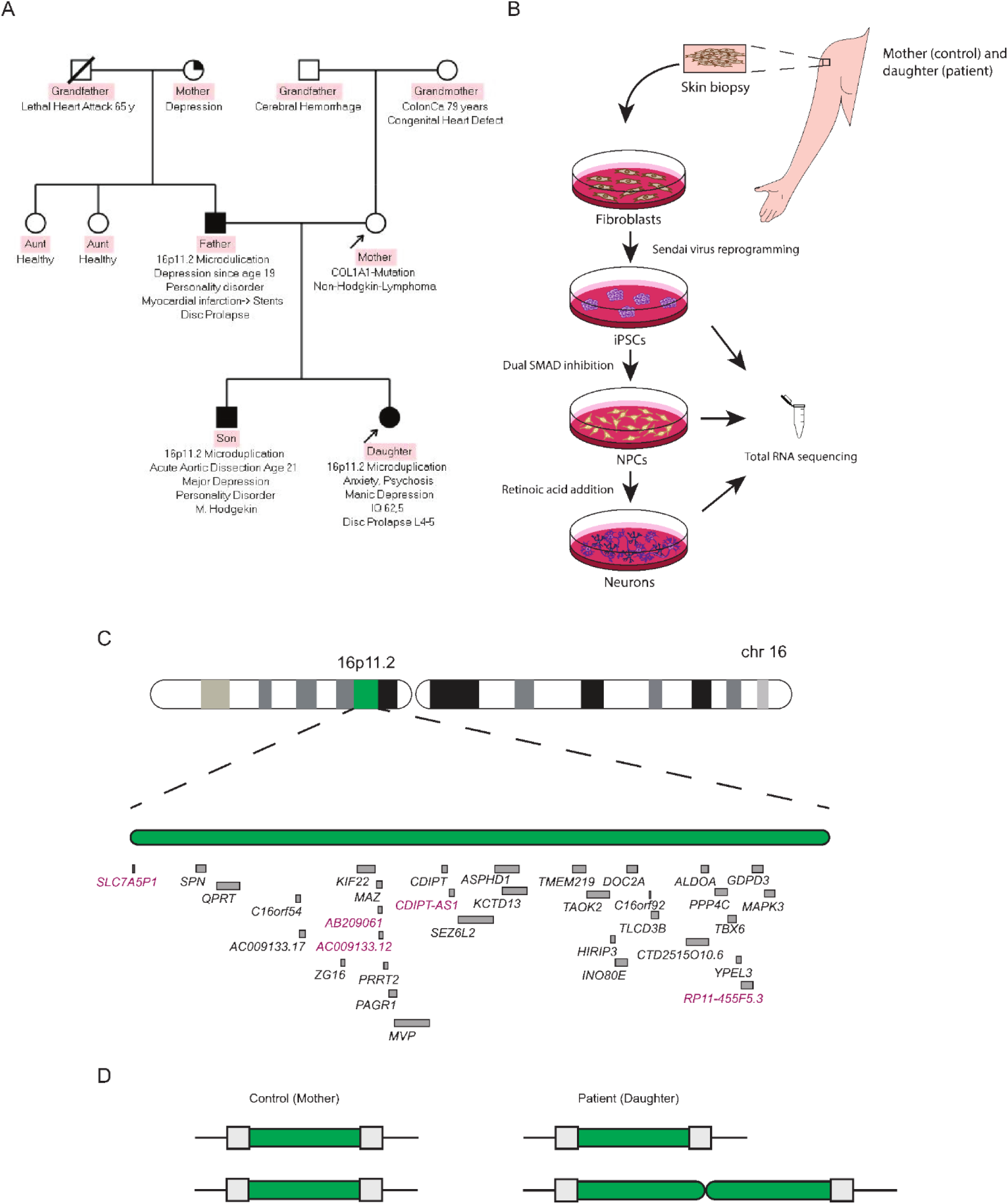
Generation of an *in vitro* neuronal model of the 16p11.2 microduplication from a female patient. A) Pedigree of a German nonconsanguineous family highlighting that the daughter, son and father are affected by an autosomal dominant 16p11.2 microduplication. Symptom summaries are provided for each affected individual. The arrows indicate the patient and control from which the skin biopsies were obtained. B) Cartoon depicting the workflow for the *in vitro* approach used. Fibroblasts were isolated from skin biopsies obtained from the non-affected mother and affected daughter and reprogrammed into iPSCs. iPSCs were then differentiated into NPCs and neurons. RNA sequencing analysis was performed by comparing iPSCs, NPCs and neurons. C) Schematic representation of the genes located in the genomic region of chromosome 16p11.2 (hg19 coordinates 29,580,020-30,176,508). Protein-coding genes are highlighted in black, whereas nonprotein-coding genes are highlighted in pink. D) Cartoon showing the heterozygous duplication of the region in the patient.

### 16p11.2 microduplication significantly altered the transcriptome of neuronal cells

To investigate the impact of the duplicated region on the phenotype associated with patients, dermal fibroblasts were isolated from skin biopsies (daughter and mother, referred to as patient and control, respectively) and reprogrammed into induced pluripotent stem cells (iPSCs). iPSCs were subsequently differentiated into a 2D neural *in vitro* model, comprising 2-step differentiation into neuronal progenitor cells (NPCs) and neurons. Successful reprogramming and differentiation were confirmed by RT-qPCR and immunostaining for iPSC, NPC, and neuronal markers (Supplemental_Fig_S1.pdf). Changes in global gene expression associated with microduplication during neurodevelopment were investigated via bulk RNA-sequencing analysis of iPSCs, NPCs, and neurons derived from the index patient and the unaffected mother. Principal component analysis (PCA) revealed that samples clustered together based on their origin (either patient or control) and cell type, confirming the cell type specificity of the RNA-seq data (Figure 2A). iPSCs from the patient and control clustered closely, indicating high transcriptomic similarity in the pluripotent state (Figure 2A). Interestingly, patient NPCs cluster relatively distant from control NPC, but rather close to control and patient neurons, while patient and control neurons are clearly separated (Figure 2A). The similarity between patient NPCs and control or patient neurons points to premature differentiation in the patient NPCs. To investigate the molecular basis of this observation, we compared expression profiles, identified overlapping genes in control neurons and patient NPCs, and performed GO term analysis (Supplemental file 1: Table_S1.xls). Eight of 40 overlapping genes were found to be associated with lipid metabolic processes, which are known to play crucial roles in neuronal differentiation, suggesting the activation of a common neural differentiation pathway in patient NPCs and control neurons (Ramosaj et al. 2021). To characterize the transcriptional changes associated with neuronal differentiation in the microduplication patient, we performed differential expression analysis comparing the patient and control iPSCs, NPCs, and neurons (Figure 2B). Overall, we observed ∼8-fold and ∼9.5-fold greater gene dysregulation (up- and downregulation relative to control) in the patient’s NPCs and neurons than in the iPSCs, respectively (Figure 2B). These findings suggest that the microduplication significantly impacted the transcriptome throughout the differentiation process, with more pronounced changes observed in the later stages of neuronal differentiation.

**Figure 2:**
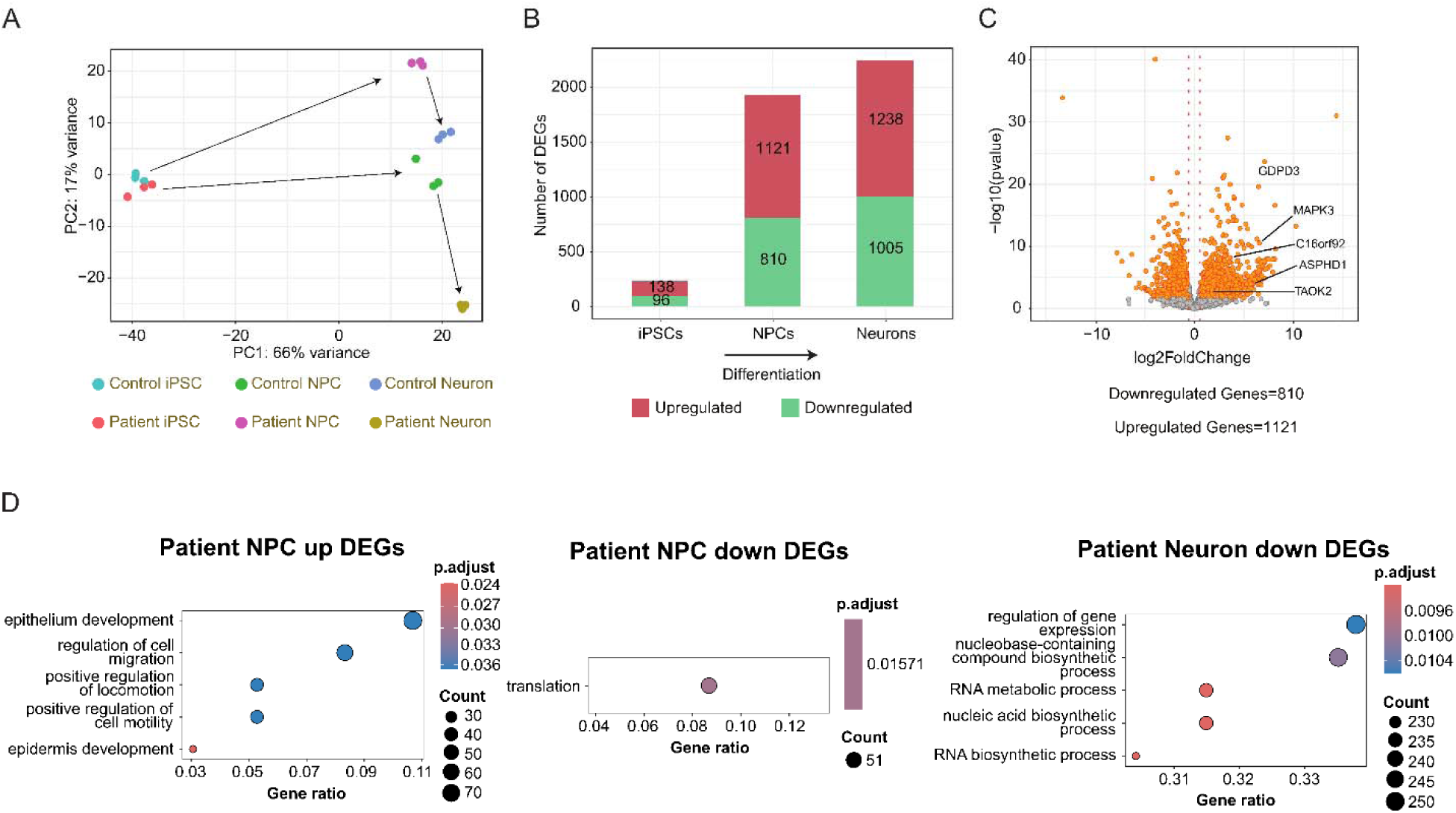
16p11.2 microduplication-dependent disruption of gene expression in neuronal cells. A) Principal component analysis (PCA) of patient and control iPSCs, NPCs, and neurons indicating transcriptomic similarities. The arrows indicate the pseudo trajectory of neural differentiation from iPSCs to NPCs and neurons. n=3 for patient and control iPSCs, NPCs, and neurons. B) Bar plot showing the number of differentially expressed genes (DEGs) between patient and control iPSCs, NPCs, and neurons. C) Volcano plot reporting the fold change of the differentially expressed genes in NPCs. The genes in orange represent significantly differentially expressed genes. Genes inside the duplicated region are highlighted. FDR <10%. Fold change: 1.5-fold. D) Gene Ontology (GO) plot representing the biological processes associated with (left) upregulated genes in patient NPCs compared with control NPCs, (middle) downregulated genes in patient NPCs compared with control NPCs and (right) downregulated genes in patient neurons compared with control neurons.

### *MAPK3* is a hub driving dysregulation in NPCs with 16p11.2 microduplication

When comparing iPSCs with NPCs and neurons, a subset of genes located in the 16p11.2 region showed substantial differences in fold-change expression during differentiation in the patient. While the expression of *MAPK3* was significantly increased in patient iPSCs (log2fold change = 2.68), in NPCs and neurons, however, *MAPK3* expression was even greater (log2fold change =6.56 in NPCs and log2fold change=3.889 in neurons). A similar expression pattern was observed for *GDPD3*. While the expression in patient IPSCs was significantly increased at an expected level (log2fold change=2.68), in NPCs and neurons, *GDPD3* upregulation was substantially greater (log2fold change=7.08 in NPCs and log2fold change=5.25 in neurons). The expression of *MAZ* did not change in the patient’s iPSCs or NPCs but showed a significant upregulation in neurons (log2fold change=0.86). Four additional genes within the microduplication region were found to be dysregulated in NPCs (Figure 2C) while showing no differential expression between the patient and control iPSCs and neurons: *ASPHD1*, *TAOK2*, *C16orf92,* and *TBX6* (log2fold change in NPCs=2.36, 1.93, 2.12, and 1.78, respectively). Even though we expected all genes within the duplicated area to be upregulated, the expression of *SLC7A5P1, SPN, QPRT, KIF22, AB209061, PRRT2, PAGR1, AC009133, MVP, CDIPT, CDIPT-AS1, SEZ6L2, KCTD13, TMEM219, HIRIP3, INO80E, DOC2A, TLCD3B, CTD2515O10.6, ALDOA, PPP4C, DGDPD3, YPEL3, and RP11-455F5.3* did not seem to be affected in iPSCs, NPCs or neurons. No genes within the duplicated region exhibited downregulation. Apart from the genes within the duplicated region, which were expected to be upregulated, genes across the entire genome were dysregulated. By intersecting the deregulated genes in the patient’s NPCs with a list of highly confident genes associated with NDD (Leblond et al. 2021), a significant overlap of 166 genes was observed (Supplemental_Fig_2.B.pdf; Supplemental file 1: Table_S2.xls), suggesting that the 16p11.2 microduplication impacts the expression of genes essential during neurodevelopment.

Gene ontology analysis of the upregulated genes in the patient’s NPCs revealed enrichment in processes related to cell motility or migration and epithelium and epidermis development (Figure 2D). For example, genes such as *MMP9, MMP14, ACTA2, LAMB2, SMAD14, KANK4*, and *VIM*, among others, regulating cell motility were found enriched in patient NPCs (Supplemental file 1: Table_S3.xls). Furthermore, a cluster of collagen-related genes, including *COL1A1, COL3A1, COL5A1, COL5A2, COL6A1, COL6A2, COL6A3, and PCOLCE*, etc., involved in collagen fibril organization processes was also found among the upregulated DEGs in the patient’s NPCs (Supplemental file 1: Table_S3.xls). Conversely, the downregulated genes showed enrichment in biological processes related to translation (Figure 2D). These processes are modulated by genes such as *MTIF2/3, EIF3A/B, EIF2AK1, RPS2/3,* among others (Supplemental file 1: Table_S4.xls). These findings suggest that the 16p11.2 microduplication leads to decreased cell proliferation and increased cell migration and motility in patient NPCs.

### Patient neurons exhibit upregulation of *MAPK3*-driven regulatory network and downregulation of chromatin remodeling genes

Gene ontology analysis of DEGs in patient neurons compared with control revealed the upregulation of key processes during neurodevelopment, such as cell-matrix adhesion, supramolecular fiber organization, dendrite extension, synaptic signaling, and chloride transmembrane transport-related processes (Supplemental file: Table_S5.xls). Similar to the NPCs, PPI network analysis highlighted *MAPK3* as a central hub that interacts with key genes involved in neurodevelopment and neuronal function such as *BDNF, PTPN7, CTNNB1, ESR1, IRS1, PTPN5, BCL2L1* and *SOS1* (Supplemental_Fig_S3.pdf). However, in neurons, we observed a milder upregulation of *MAPK3* in the patient compared to the control (log2foldchange = 3.89), suggesting a less pronounced *MAPK3*-related effect in neurons than in NPCs.On the other hand, genes downregulated in the patient’s neurons were enriched for terms related to regulation of gene expression and RNA metabolic processes (Figure 2D). These include chromatin remodelers such as *CHD7* and *BRD9*, SWI/SNF-related actin-dependent chromatin regulators (*BCL7A, SMARCE1, SMARCB1, and SMARCC1*), the histone deacetylase *HDAC1* and transcription factors (*PHF10, TBP, FOXP1, and SOX11*), all of which contribute to chromatin remodeling and transcriptional regulation (Supplemental file 1: Table_S6.xls). In summary, similar to findings in NPCs, our analysis of patient and control neurons identified *MAPK3* as a regulatory hub driving gene upregulation in differentiating neurons carrying the 16p11.2 microduplication.

### Distinct transcriptional profiles in NPCs and neurons highlight key genes affected by chromosome 16 microdeletion and microduplication

To complement the analysis performed on the neuronal cells of a patient carrying a 16p11.2 microduplication, we compared our RNA-seq data with RNA-seq data of NPCs and neurons of two patients with a 16p11.2 microdeletion generated by Liu F. and colleagues (Liu et al. 2023). Similar to the patients with microduplications, patients with microdeletions present with dysmorphic features, behavior abnormalities, and psychiatric symptoms. However, while several microduplication cases are accompanied by microcephaly, microdeletion cases are characterized by macrocephaly (Auwerx et al. 2024a; Elsayed et al. 2024; McRae et al. 2025). A comparison of DEGs revealed genes whose expression was upregulated in the microduplication and down-regulated in the microdeletion NPCs and neurons (Figure 3A). Overall, 29 such genes, including *MAPK3, TAOK2, GDPD3, C16orf92*, and *ASPHD,* were identified in NPCs. A network analysis of the 29 genes confirmed MAPK3 as a regulatory hub with protein-protein interactions linking it to six other genes, four of which reside within the duplicated/deleted region (Figure 3B). Comparison of the expression dynamics of genes in the affected 16p11.2 segment revealed that *MAPK3, ASPHD1, C16ORF92, GDPD3, MAZ*, and *TAOK2* showed mostly opposite expression profiles in deletion (blue dots) and duplication (green dots) conditions in NPCs and neurons (Figure 3C). *MAZ* appeared to be upregulated only in neurons but not in NPCs. The rest of the genes within the 16p11.2 region seemed unaffected by duplication/deletion or presented expression patterns that deviated from dosage-dependent trends. In neurons, a total of 7 genes, including *GDPD3*, *MAZ*, and *MAPK3* from the affected chromosomal segment, were upregulated in the microduplication group and downregulated in the microdeletion group (Figure 3D). Overall, the comparison of the transcriptional changes caused by microduplication and microdeletion confirmed that a subset of genes exhibited reciprocal expression patterns under these conditions and that *MAPK3* was a key regulator driving the dysregulation of several genes from inside and outside the affected genomic segment on chromosome 16.

**Figure 3:**
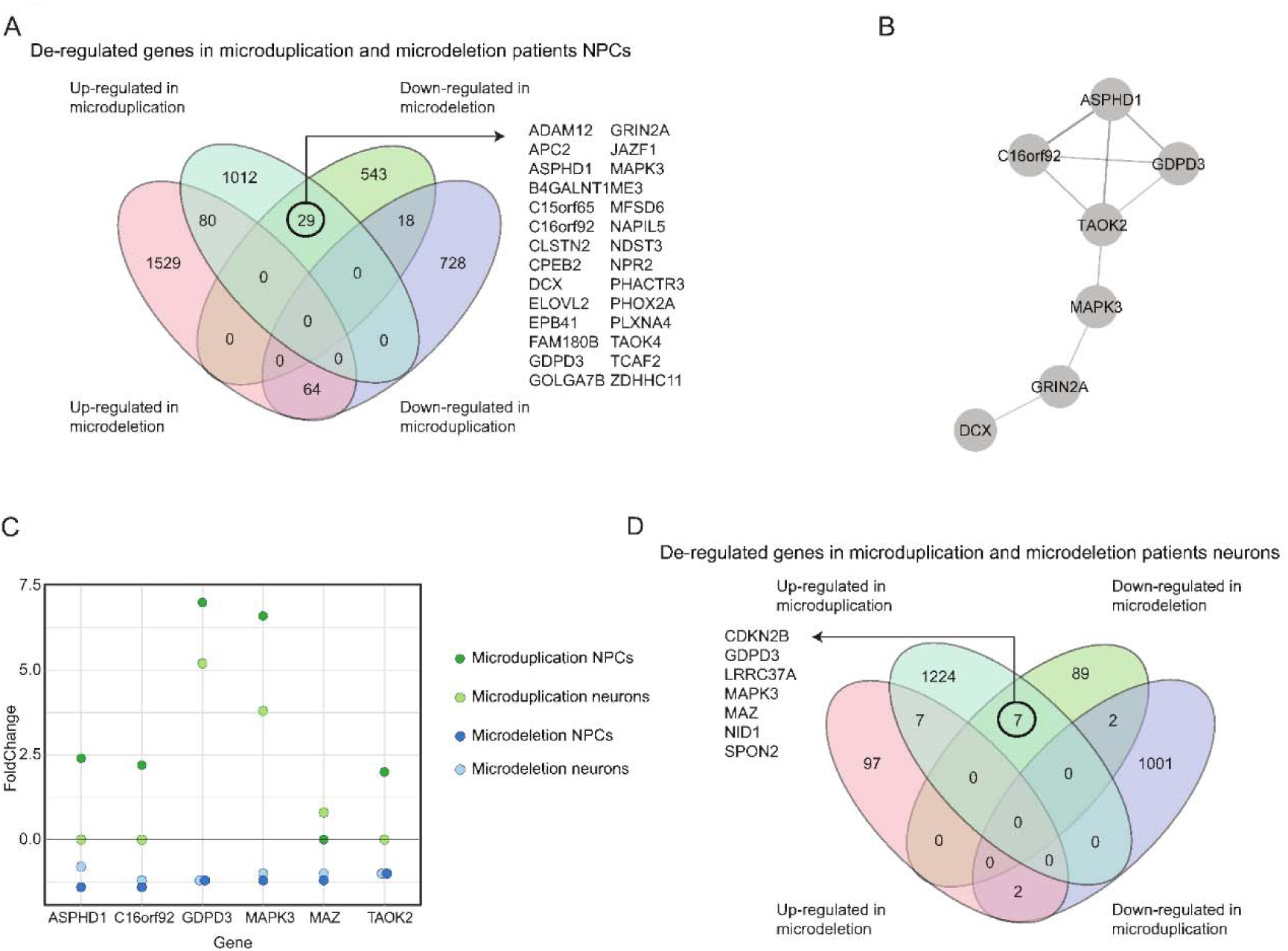
Comparison of the deregulated genes in the 16p11.2 microduplication and microdeletion revealed MAPK3 as a central player in the disease phenotype. A) Overlap of deregulated genes in NPC from 16p11.2 microduplication and 16p11.2 microdeletion patients compared with controls. Genes inside the 16p11.2 region are highlighted in red. B) Protein-protein interaction network for upregulated genes in the microduplication and downregulated in the microdeletion carrying cells. C) Plot reporting the deregulation of genes located within the 16p11.2 region in NPCs and neurons derived from patients with microduplication (green dots) and microdeletion (blue dots). D) Overlap of deregulated genes in neurons from 16p11.2 microduplication and 16p11.2 microdeletion patients compared with controls. Genes inside the 16p11.2 region are highlighted in red.

### Intrachromosomal gene clusters and TAD-associated co-regulation of differentially expressed genes

In addition to transcriptional changes, CNVs can influence higher-order chromatin organization, altering regulatory contacts and thereby contribute to transcriptional dysregulation. We therefore asked whether duplication of the 16p11.2 region is associated with altered chromatin contacts either locally, within the duplicated locus, or more broadly across the genome. We generated Hi-C data from patient and control iPSCs and NPCs and first examined chromatin interactions across the 16p11.2 region (Figure 4A). In patient-derived iPSCs, we identified a region of significantly increased chromatin contacts within the duplicated interval. This local increase was not observed in NPCs, indicating that the chromatin effects of the duplication may differ between cellular states. We then investigated whether genome-wide compartment changes were associated with transcriptional alterations. Differential compartments were separated according to direction of change and intersected with up- and downregulated DEGs. This analysis showed the clearest overlap between increased compartments and upregulated DEGs, particularly in NPCs (Figure 4B). In contrast, decreased compartments showed limited overlap with downregulated DEGs. The absence of a strong signal in iPSCs may be partly explained by the smaller total number of DEGs detected in this cell type. The generally weaker association between decreased compartments and downregulated genes suggests that loss of compartment activity may not necessarily be sufficient to reduce expression, possibly because remaining chromatin contacts sufficiently continue to support transcription. Overall, these results suggest that altered chromatin organization may contribute to gene expression changes associated with 16p11.2 duplication, especially in NPCs.

**Figure 4:**
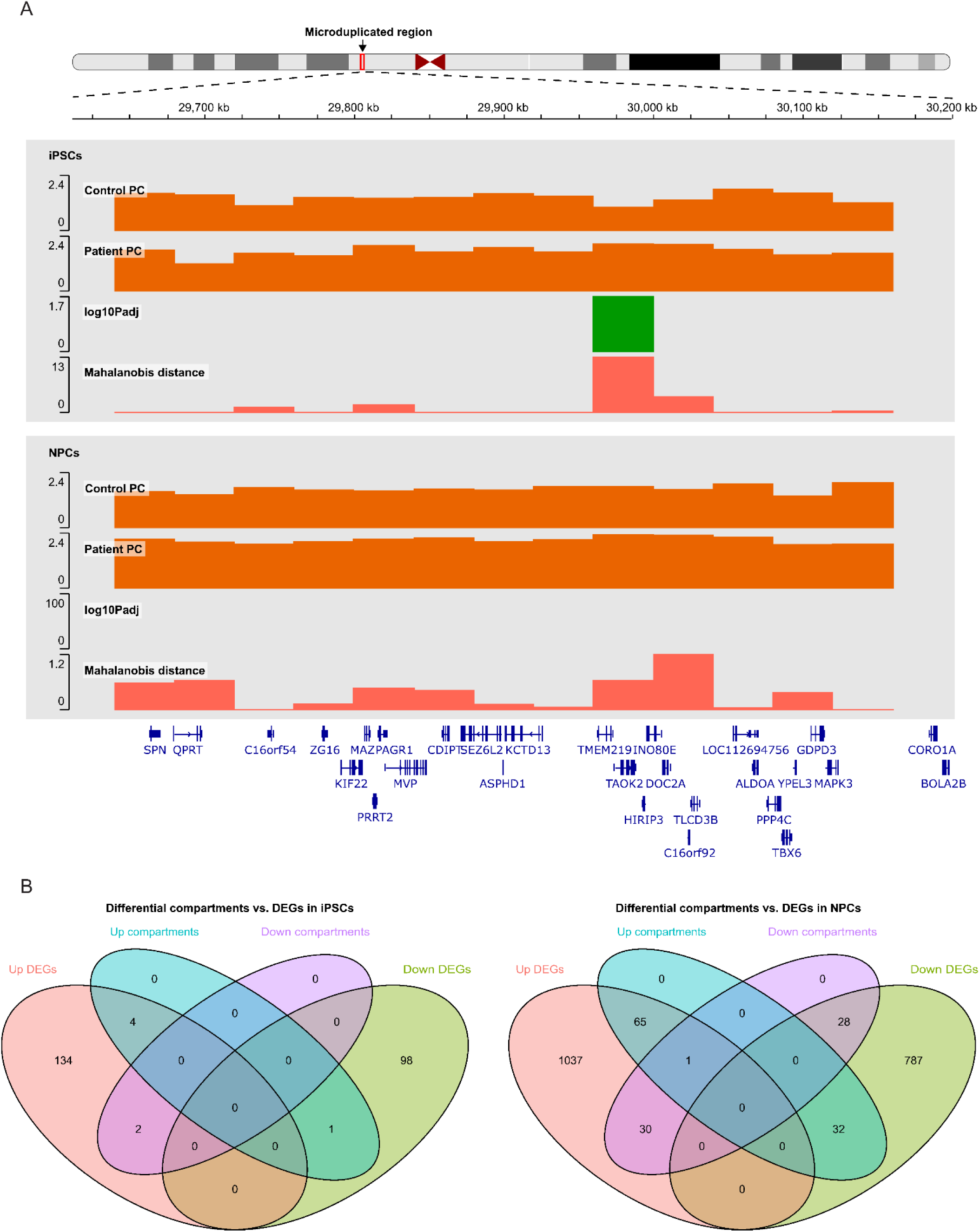
16p11.2 microduplication-associated DEGs co-occur with differential chromatin compartments. A) Differential chromatin compartment analysis across the 16p11.2 region in iPSCs and NPCs. Principal component values derived from Hi-C compartment analysis are shown for control and patient cells in each genomic bin, together with the corresponding adjusted p value and Mahalanobis distance used to identify differential compartment bins. B) Overlap between differential chromatin compartment bins and differentially expressed genes. Increased and decreased compartment bins were intersected separately with upregulated and downregulated DEGs in iPSCs and NPCs.

## Discussion

Copy number variations (CNVs) play essential roles in genome dynamics and can be key contributors to disease pathogenesis. While the influence of dosage-sensitive genes within CNVs on disease mechanisms is widely described and accepted, the impact of the noncoding genome and its functional contribution remain largely elusive (Pos et al. 2021). In line with this, the mechanisms underlying the tissue-specific pathological effects of CNVs are not clearly understood. Although CNVs often affect multiple genes at the genomic level, microdeletion or microduplication syndromes can result in highly specific phenotypes with restricted and very selective tissue involvement (Shaikh 2017). The most obvious explanation is the tissue-specific expression of the affected genes. However, since broadly expressed genes are often affected, additional mechanisms must be taken into account. Tissue-specific interaction partners, vulnerability during narrow developmental time windows, and selective functionality in a specific temporal and spatial context are further and more complex factors that must be considered.

By studying a family carrying a 16p11.2 microduplication that segregates with a variety of psychiatric aberrations over two generations, we provide evidence that neuronal differentiation significantly increases the impact of the microduplication on gene expression. This effect involves additional genes both within and beyond the critical region and leads to an increase in the fold change in expression between patient and control far beyond a 2.0 score, which would typically be expected in the context of a microduplication. By searching for potential hubs, we identified a network centered around MAPK3, which, through enhancement of the MAPK/ERK pathway, contributes to the observed gene expression alterations. However, in addition to this interaction network, we identified a substantial number of unrelated genes that were significantly up- or downregulated in the patient’s cells. Several of these genes cluster within TADs overlapping the 16p11.2 region, particularly in NPCs and neurons, suggesting that a pathological chromatin architecture may contribute to the dysregulated gene expression observed in microduplication 16p11.2 carriers during neurodevelopment. Analysis of comparable transcriptome data of patients with the corresponding 16p11.2 microdeletion also revealed a MAPK3-centered network of genes, suggesting reciprocal expression behavior of several MAPK3-related genes in microdeletion/microduplication 16p11.2 patients. Unlike in microduplication-carrying NPCs and neurons, genes unrelated to the MAPK3 hub in microdeletion data did not cluster at the chromatin interaction sites of 16p11.2. Only the microdeletion itself formed a coregulated gene cluster.

Genomic rearrangements at 16p11.2 constitute one of the most frequent causes of genomic disorders. Both 16p11.2 microdeletions and microduplications contribute to a broad pleiotropic spectrum of phenotypic traits, including neurodevelopmental and neuropsychiatric diseases (Hudac et al. 2020; Rein and Yan 2020; Auwerx et al. 2024b; McRae et al. 2025). While 16p11.2 CNVs show high variability in expressivity and reduced penetrance, some reciprocal features are found in patients, including microcephaly and a lower body mass index (BMI) in microduplication patients and macrocephaly and obesity in microdeletion patients (Qureshi et al. 2014; Auwerx et al. 2024a). In line with such partially reciprocal phenotypes, we here found MAPK3 as a key regulator of gene expression changes in differentiating neuronal cells harboring either the microduplication or the microdeletion. Together with *MAPK3,* 28 other genes, including *ASPHD1*, *C16orf92*, *GDPD3* and *TAOK4,* are downregulated in NPCs from patients with microdeletion, and only *ASPHD1*, *C16orf92*, *GDPD3* and *TAOK4* are upregulated in the microduplication patient. In neurons, seven genes, including *MAPK3* and *GDPD3,* are downregulated in microdeletion carriers, whereas only *MAPK3* and *GDPD3* are upregulated in the microduplication carrier. The contrasting expression response of these genes to the microduplication/microdeletion 16p11.2 suggests their crucial role in driving the reciprocal phenotypic features of the two syndromes.

MAPK3 seems to play a critical role in driving pathology in both 16p11.2 microduplication and microdeletion syndromes. *MAPK3* (also known as *ERK1*) (Roskoski 2012), which emerged as a key player in our analysis, was significantly upregulated in the microduplication patient’s and downregulated in the microdeletion patient’s NPCs and neurons. It is a component of the mitogen-activated protein kinase (MAPK) signaling pathway and plays a critical role in transducing extracellular signals to the nucleus to regulate gene expression. MAPK3 is an important driver of developmental disorders, particularly rare disorders called RASopathies, which are characterized by a broad spectrum of neuropsychiatric features, including intellectual disabilities and autism spectrum disorders. Its role in neurogenesis also points toward a function in the regulation of brain growth, supporting its involvement in macro- and microcephaly. It has been suggested that, among other factors, altered ERK1 signaling during critical stages of brain development disrupts synaptic connectivity and neuronal function (Lavoie et al. 2020; Iroegbu et al. 2021).

The upregulation of ERK1 has also been linked to autism (Yang et al. 2011; Zou et al. 2011), underscoring the intricate balance required for proper ERK1 function in maintaining neuronal homeostasis. Our PPI networks revealed connections between MAPK3 and key regulators of cellular functions such as BDNF, MMP9, CASP8, PXDN, PLCG1, and IRS1. Phospholipase C gamma (PLCG1) is an enzyme involved in transducing signals from cell surface receptors to intracellular signaling pathways and can influence the MAPK3 pathway through downstream effects (Diaz-Flores et al. 2013). Similarly, IRS1 activates the MAPK3 pathway through insulin signaling, facilitating the Grb2-SOS-Ras pathway and contributing to cellular differentiation and proliferation (Boucher et al. 2014; Rabiee et al. 2018). Furthermore, MAPK3 regulates critical genes such as BDNF and MMP9 (Kuzniewska et al. 2013). BDNF (brain-derived neurotrophic factor) is an important regulator of synaptic transmission and long-term potentiation, thereby influencing memory processes (Leal et al. 2014). MMP9 is involved in extracellular matrix remodeling and, through this process, is an important regulator of neurodevelopmental processes (Reinhard et al. 2015). Similarly, Peroxidasin (PXDN) contributes to extracellular matrix remodeling, promoting cell migration and angiogenesis (Medfai et al. 2019; Lee et al. 2020). Caspase-8 (CASP8) plays a key role in cell adhesion processes (Finlay and Vuori 2007). Overall, our results indicate that MAPK3 modulates the regulation of critical genes such as *BDNF*, *MMP9*, and *CASP8,* influencing processes such as extracellular matrix assembly, collagen fibril organization, and actin filament dynamics, with potential implications for neurodevelopmental disorders.

The significant role of MAPK3 and its downstream targets in the transcriptional aberrations of differentiating neural cells in a 16p11.2 microduplication carrier suggests that MAPK3 inhibition could be a potential therapeutic approach for patients with this CNV. To date, no MAPK3 inhibitor therapies have received clinical approval. However, natural anthraquinones such as emodin-8-glucoside, as well as synthetic ERK inhibitors such as LY3214996, have shown promising results in preclinical studies for the treatment of cancers driven by the ERK pathway (Bhagwat et al. 2020; Jamshidi et al. 2024). Nonetheless, evidence supporting the use of MAPK3 inhibitors in neurodevelopmental disorders remains limited. In a study by Blizinsky et al. (Blizinsky et al. 2016), pyramidal neurons carrying a 16p11.2 microduplication presented greater dendritic complexity than did wild-type neurons. When the cells were treated with an ERK inhibitor during incubation, a reduction in dendritic complexity was observed, suggesting a potentially successful therapeutic route for patients with the microduplication 16p11.2.

Surprisingly, we found a substantial increase in the fold changes of DEGs beyond expectations during the neuronal differentiation of microduplication patient cells, particularly for genes within the critical region. Some of the DEGs interact within a network centered around MAPK3, suggesting a cumulative gain-of-function effect. This effect may amplify changes in gene expression throughout the network because of the role of MAPK3 in signal transduction. Additionally, the microduplication may alter chromatin structure, thereby altering chromatin interactions in the 16p11.2 region and increasing the susceptibility of genes within the critical region to chromatin-mediated expression changes. During neuronal differentiation, particularly when stem cells differentiate into neuronal precursor cells, massive remodeling of chromatin takes place (summarized in (Decker et al. 2020; Martino et al. 2023)). During this remodeling process, chromatin interactions of 16p11.2 might follow different paths in a wildtype compared to a microduplication configuration. This could, in turn, place 16p11.2 under the influence of active chromatin marks, leading to a disproportionately high increase in expression-far beyond what would be expected on the basis of genomic duplication alone.

A large proportion of the DEGs in our dataset, however, are not located in the critical region and are not connected through gene-gene interactions with genes within the critical region. Structural aberrations have been shown to interfere with the 3D structure of chromatin, thereby influencing gene expression (Rovina et al. 2020; Chilinski et al. 2022; Franke et al. 2022). Here, we hypothesized that chromatin dynamics during neuronal differentiation are particularly vulnerable to aberrations in the DNA polymer, rendering them susceptible to conformational changes induced by CNVs. This not only leads to dramatic changes in the expression of genes within the critical region but also extends beyond it, influencing chromatin regions that interact with 16p11.2 through large-scale compartmental interactions (Dixon et al. 2012; Papantonis and Cook 2013; Smith et al. 2016). To directly investigate whether the 16p11.2 microduplication is accompanied by changes in 3D genome organization, we generated Hi-C data from patient and control iPSCs and NPCs. At the local level, this analysis identified increased chromatin contacts within the duplicated 16p11.2 region in patient-derived iPSCs. This effect was not detected in NPCs, suggesting that the impact of the duplication on local chromatin architecture may be cell-state dependent. We next examined whether broader chromatin compartment changes were linked to transcriptional alterations. Genome-wide differential compartment analysis revealed that increased compartment bins preferentially overlapped with upregulated DEGs, with the clearest association observed in NPCs. By contrast, decreased compartment bins showed limited overlap with downregulated DEGs. These findings suggest that increased compartment activity may be associated with transcriptional activation in the context of 16p11.2 microduplication, particularly during neural progenitor stages. More broadly, our Hi-C analysis supports a model in which CNV-associated changes in chromatin organization may modulate the transcriptional consequences of the duplication beyond local gene dosage effects. In view of the extensive chromatin remodeling that occurs during neurodevelopment, these observations are consistent with the idea that the nervous system may be particularly vulnerable to CNVs that perturb genome organization and regulatory interactions, making it the organ that is most often and significantly affected by CNVs (Rein and Yan 2020). On the other hand, the expression of genes within the critical region was halved in iPSCs, NPCs and neurons derived from microdeletion patients, as expected on the basis of their genomic CNV. A detailed SGC analysis of the differentially expressed genes unrelated to the genes within the critical region and a comparison with established TADs in NPCs and neurons, other than in the cells carrying the microduplication, revealed only one SGC corresponding to the 16p11.2 region but in no other part of the genome. This observation suggests that, unlike the 16p11.2 microduplication, the microdeletion has a minimal effect on chromatin remodeling during neurodevelopmental trajectories. Further studies using *in vitro* and animal models are needed to better understand chromatin remodeling processes during neuronal differentiation and their susceptibility to chromatin aberrations.

In summary, we analyzed the neuronal molecular phenotype of a family harboring a 16p11.2 microduplication via an *in vitro* model of neurodevelopment. We leveraged genome-wide transcriptomic studies and complemented our findings with a reanalysis of published microdeletion datasets and newly generated Hi-C data. Our analysis identified critical time points during neuronal differentiation at which gene expression changes exceeded expectations on the basis of the 16p11.2 microduplication alone. Furthermore, we highlighted MAPK3-centered gene hubs among the dysregulated genes in both microduplication- and microdeletion-carrying cells, suggesting that MAPK3 is a potential therapeutic target for 16p11.2 CNV patients. Finally, we found evidence that changes in chromatin conformation substantially contribute to disease-related transcriptional alterations in microduplication cells during neuronal differentiation, supporting the idea that the structural genome context can play a key role in modulating the impact of CNVs.

## Methods

### Ethical approval

The study was approved by the Landesärztekammer Rheinland-Pfalz for the study of patient-derived iPSCs as a model for neurogenetic diseases (committee’s reference number for approval: 837.156.15 (9924)/2015). The proband, her brother and both parents consented to the research study and provided written informed consent for participation.

### Chromosome Microarray Analysis

Blood was collected from the proband, her brother and parents, and chromosome microarray analysis (CMA) was performed via the Affymetrix CytoScan® HD array (Affymetrix Inc., Santa Clara, CA, USA) according to the manufacturer’s instructions. This platform contains a total of almost 2.7 million copy number markers (1,953,246 CN probes + 743,304 SNP probes) with a 0.88 kb mean probe spacing with respect to RefSeq genes. The data were analyzed via Chromosome Analysis Suite software version 3.1.1.27 (Affymetrix) at a resolution of 100 kb. In addition, deletions were analyzed at higher resolution (5 markers per 1 kb). Identifications of CNVs were compared with the following public databases: the DECIPHER database (http://decipher.sanger.ac.uk/), the Database of Genomic Variants (http://dgv.tcag.ca/dgv/app/home), OMIM (http://www.omim.org), UCSC (http://genome.ucsc.edu/, hg19), the International Standards for Cytogenomic Arrays (https://www.iscaconsortium.org/) and the Ensembl database (https://www.ensembl.org).

### Fibroblast isolation

Four-millimeter skin biopsies were taken at the Children’s Hospital of the University Medical Center in Mainz. Fibroblasts were isolated as previously described with small adaptations (Vangipuram et al. 2013). In summary, skin biopsies were cut into 18-24 equally sized pieces containing all the skin layers. The pieces were plated on a six-well plate that was previously coated with 0.1% gelatin. Fresh media was added every other day. Fibroblasts started to migrate out of the skin biopsies after 7-10 days and were split into two 75cm^2^ flasks after 3-4 weeks. When 90% confluence was reached, the two flasks were split into 3 175cm^2^ flasks. Fibroblasts could be expanded as needed or frozen in liquid nitrogen for long-term storage and later experiments. Cells were routinely screened for mycoplasma contamination.

### iPSC reprogramming and culture

Fibroblasts were reprogrammed into iPSCs by following the feeder-dependent protocol of the CytoTune(TM)-iPS 2.0 Sendai Reprogramming Kit (Thermo Fisher Scientific) with small adaptations. Briefly, on day 0, fibroblasts were transduced with Sendai virus-derived vectors containing the four Yamanaka factors OSKM (Oct3/4, SOX2, Klf4, and c-Myc). After seven days, the cells were split into mouse embryonic fibroblast (MEF)-feeder layer cells, and one day later, the medium was changed to iPSC medium. After three to four weeks, iPS colonies started to emerge, and 40-50 colonies were manually picked and transferred to one well of a Matrigel (Coarning)-coated 12-well plate. The medium was changed to mTeSR1 (Stem Cell), and the cells were expanded until ready for characterization. For the removal of the remaining viral particles, the cells were cultured for 5-7 days at 39°C. iPSCs were routinely cultured in mTeSR1 on Matrigel-coated 6-well plates in a humidified incubator at 37°C and 5% CO2. Cells were typically split every 5-7 days at a ratio of 1:3 to 1:10 using custom-made enzyme-free splitting buffer containing PBS, NaCl and EDTA. Cells were routinely tested for mycoplasma contamination. A schematic representation of the workflow of the generation of iPSCs is shown in **Figure 1**.

### Karyotyping

After stable and virus-free culture was achieved, iPSCs were seeded into 25cm^2^ flasks coated with Matrigel. The cells were grown until nearly ready to split. For metaphase arrest, colcemid was added to the cells, which were then incubated for 60 minutes at 37°C. The cells were subsequently harvested and treated twice with a hypotonic KCl solution and fixed with a 1:3 and 1:2 solution of methanol and ethanoic acid. The cell suspension was then dripped on microscope slides and dried on a 42°C hot plate. After drying, the cells were incubated at 200°C for 1-2 hours, stained with quinacrine, embedded and analyzed via a nonconfocal fluorescence microscope and Applied Biosystems imaging software.

### iPSC differentiation into NPCs

iPSC differentiation into NPCs was performed by using PSC Neural Induction Medium (NIM, Gibco). On day −1, iPSCs were seeded on a Matrigel-coated 6-well plate. After 24 hours, for the cells that reached a confluency of 15-25%, the medium was changed to neural induction medium (NIM: Neurobasal (Thermo Fisher), 2% neural induction supplement (Thermo Fisher), and 1% P/S), and medium was replaced every other day. On day 7 of neural induction, the cells were replated following the manufacturer’s protocol in a Geltrex-coated (Thermo Scientific) 6-well plate (500×103 cells/well) in Neural Expansion Medium (NEM: 49% Neurobasal, 49% Advanced DMEM (Thermo Scientific), 2% Neural Induction Supplement, 1% P/S) supplemented with ROCK inhibitor (5 μM, Stem Cell). After 24 hours, the medium was exchanged with NEM without ROCK inhibitor. The cells were monitored daily, and the medium was changed every other day. When the cells reached confluency, they were replated via TrypLE™ Express and cultured in poly-ornithine/laminin-coated dishes with NM supplemented with FGF2 (20 ng/ml). Cells routinely reported negative for mycoplasma contamination. A schematic representation of the workflow of NPC differentiation is shown in **Figure 1**.

### NPC differentiation into neurons

For neuronal differentiation, NPCs were plated at low density (50-100 x10^3^ cells/well) on poly-ornithine/laminin-coated 6-well plates. Twenty-four hours after seeding, the cells were washed twice with PBS and cultured with NM+VitA medium (DMEM/F-12, 1% N2 supplement, 2% B27+VitA supplement (Thermo Fisher), 1% P/S). 8% CO2 was used for culturing the cells. Fresh medium was added to the cells every 3-4 days. When a total volume of 10-12 ml per well was reached, 20-50% of the medium was removed before fresh media was added. After 35 days, differentiated neurons were harvested as needed. Cells were routinely screened for mycoplasma contamination. A schematic representation of the workflow of neuronal differentiation is shown in **Figure 1**.

### Immunofluorescence staining

For immunofluorescence staining, the cells were seeded on 18 mm coverslips and cultured for 2-3 days. All the following steps were conducted at room temperature. The cells were fixed for 20 minutes in 4% PFA in PBS. The cells were washed three times with PBS, followed by a 30-60-minute incubation with blocking solution (5% BSA (company) and 0.3% Triton X-100 (Carl Roth) in PBS). Afterwards, the cells were incubated with primary antibodies diluted in blocking solution at +4°C overnight (ON). The next day, the cells were again washed three times with PBS containing 0.1% Triton X-100 and incubated for one hour with secondary antibodies (1:800) diluted in blocking solution in the dark. Three washing steps with 0.3% Triton X-100 in PBS followed, and coverslips were mounted on microscope slides with Vectashield (Company) containing 0.5% DAPI for nuclear counterstaining. Pictures were acquired via a confocal laser scanning microscope.

The following antibodies were used: rabbit anti-SERPHIN1 (Sigma-Aldrich; S5950; 1:500), goat anti-NANOG (R&D Systems; AF1997; 1:200), mouse anti-TRA1-60 (Millipore; 4360; 1:200), rabbit anti-PAX6 (Biolegend; 901301; 1:50), rabbit anti-SOX2 (Abcam; ab137385; 1:300), mouse anti-NESTIN (Merck; MAB5326; 1:200), mouse (IgG2b) anti-TUBB3 (Sigma; T8660; 1:300), mouse (IgG1) anti-MAP2 (Sigma-Aldrich; M4403; 1:300), and rabbit anti-TAU (Abcam; ab32057; 1:200). The secondary antibodies used were as follows: goat anti-mouse IgG Alexa Fluor 488 (Invitrogen; A11017; 1:500), goat anti-rabbit IgG Alexa Fluor 488 (Invitrogen; A11008; 1:500), goat anti-rabbit IgG Alexa Fluor 596 (Invitrogen; A11012; 1:500), rabbit anti-goat IgG Alexa Fluor 568 (Invitrogen; A11079; 1:500), and rabbit anti-mouse IgG Alexa Fluor 488 (Invitrogen; A11059; 1:500).

### RT-qPCR

Total RNA was isolated from the cell pellet via the High Pure RNA Isolation Kit (Roche). Five hundred nanograms of RNA were reverse transcribed into cDNA using PrimeScript (TM) RT Master Mix (Takara). qPCR was conducted on an Applied Biosystems StepOne Plus(TM) real-time PCR system with SYBR(R) Premix Ex Taq(TM) II and ROX plus (Takara) by using different pairs of primers (Supplemental file 1: Table_S17.xls) that were manually designed and tested. The expression levels were quantified in comparison with those of GAPDH.

### RNA-seq

NGS library preparation was performed with Illumina Stranded Total RNA Prep Ligation with Ribo-Zero Plus Kit following Stranded Total RNA Prep Ligation with Ribo-Zero Plus ReferenceGuide (Document # 1000000124514 v02 April 2021). Libraries were prepared with a starting amount of 1000ng and amplified in 9 PCR cycles. Two post-PCR purification steps were performed to exclude residual primer and adapter dimers. Libraries were profiled in a DNA 1000 chip on a 2100 Bioanalyzer (Agilent technologies) and quantified using the Qubit dsDNA HS Assay Kit, in a Qubit 4.0 Fluorometer (Life technologies). All 18 samples were pooled in equimolar ratio and sequenced on 2 NextSeq500 Highoutput FC, PE for 2x 78 cycles plus 10 cycles for the index read.

### Hi-C seq

Bulk Hi-C library preparation was performed as previously described (https://www.nature.com/articles/s41586-023-05794-2). Briefly, 2 million cells were collected and fixed with 2% formaldehyde and reaction was quenched with glycine. Cells were lysed and resulting nuclei were subjected to MboI restriction enzyme digestion, followed by biotinylation of digested ends. Next, proximity ligation was performed to connect the 3D interactions into linear fragments. Chromatin was decrosslinked and DNA was extracted via ethanol precipitation. Resulting DNA was sonicated using a Covaris E220 focused-ultrasonicator and double-side size selected using AMPure XP beads (Beckman Coulter) to obtain fragments of 300-700bps. Biotinylated DNA was pulled down on Dynabeads MyOne Streptavidin C1 (Thermo Fisher Scientific). On the bead-bound DNA fraction, end polishing, polyA tailing, and Illumina TruSeq unique dual indices (IDT) ligation reactions were performed. Lastly, libraries were amplified via PCR for 6 cycles and sequenced on an Illumina NextSeq 2000 system with a PE150 configuration.

### RNA-seq analysis

Raw data quality was checked via FASTQC v0.10.5 (https://www.bioinformatics.babraham.ac.uk/projects/fastqc). Samples passing the quality threshold were aligned to the hg19 genome via STAR v2.15a (https://github.com/alexdobin/STAR) using paired-end mode, and output was generated as coordinate-sorted BAM files. Mapped reads were quantified at the gene level with strand-specific using HTSeq-count (https://htseq.readthedocs.io/). Gene-level count matrices were then imported into DESeq2 (https://bioconductor.org/packages/release/bioc/html/DESeq2.html) for normalization and differential expression analysis in R. Statistical significance was assessed using a false discovery rate (FDR) threshold of 0.1. Variability between patient and control cell types was assessed by principal component analysis (PCA) in R using prcomp on the 500 most variable genes. Genes were defined as differentially expressed if they met the following criteria: absolute log2 fold change > 0.58, nominal p value < 0.05, and false discovery rate (FDR) < 0.1. Gene Ontology (GO) enrichment analysis was performed in R using the clusterProfiler package with the org.Hs.eg.db annotation database. Differentially expressed genes were separated into upregulated and downregulated sets and analyzed independently for enrichment of Biological Process terms using a Benjamini-Hochberg-adjusted significance cutoff of 0.1 using the full tested gene set as background. Protein-protein interaction (PPI) networks were predicted using the STRING database, integrating evidence from experimental data, co-expression, genomic neighborhood, curated databases, and text mining. Interactions with a confidence score ≥ 0.8 and node degree ≥ 5 were retained to identify the most likely interaction network. The resulting PPI network was imported and visualized in Cytoscape v3.8.0.

### Hi-C analysis

Hi-C sequencing data were analyzed using the nf-core/hic pipeline v2.1.0 (https://nf-co.re/hic/2.1.0/), using GRCh38 as the reference genome and MboI as the restriction enzyme. Contact maps were generated at 40 kb bin resolutions. Differential chromatin compartments were then called and visualized using dcHiC v2.1 (https://github.com/ay-lab/dcHiC). Directional compartment changes were assigned from dcHiC’s quantile-normalized compartment scores by computing, for each 40 kb bin, the difference between the condition-mean PC1 values (ΔPC1 = PC1_patient - PC1_control); bins with ΔPC1 > 0 were classified as “up” and bins with ΔPC1 < 0 as “down”, and these sets were used for downstream analyses.

### Data access

The datasets generated and/or analyzed during the current study will be made publicly available upon publication, due to the sensitivity of the human data. Prior to publication, data are available from the corresponding author upon reviewers’ request. Differentially expressed genes for microdeletion NPCs and neurons were obtained from Liu et al, 2023.

### Competing interest statement

The authors declare that the research was conducted in the absence of any commercial or financial relationships that could be construed as potential conflicts of interest.

### Ethics statement

This study adheres to the principles of the Helsinki Declaration. Ethical approval was obtained from the ethics committee of the Landesärztekammer Rheinland-Pfalz for the study of patient-derived iPSCs as a model for neurogenetic diseases (committee’s reference number for approval: 837.156.15 (9924)/2015). The proband, her parents and her brother consented to the research study and provided written informed consent for participation.

### Consent for publication

The proband, her parents and her brother provided written informed consent for genetic testing and publication of clinical and genetic data according to the German bioethics laws.

## Acknowledgments

The authors are grateful for the family’s cooperation in the diagnostic process and further analyses. We also thank the diagnostic and research technicians in the laboratory of the Institute of Human Genetics Mainz for their excellent technical assistance. Support by the IMB Genomics Core Facility and the use of its NextSeq500 (funded by the Deutsche Forschungsgemeinschaft (DFG, German Research Foundation) – INST 247/870-1 FUGG) is gratefully acknowledged.

## Funding

SuS and S.G. acknowledge funding by SFB 1551 Project No. 464588647 of the Deutsche Forschungsgemeinschaft (DFG). SuS acknowledges funding from the Forschungsinitiative Rheinland-Pfalz and the ReALity initiative of the Johannes Gutenberg University Mainz.

## Authorś contributions

SuS, DP and JF conceived the study, and SuS, SG and MB planned and supervised the experimental work. SuS, DP, MD, KK, OT, and SD were involved in the clinical diagnosis, management and treatment of the family. ML and JW carried out the microarray analyses and generated the clinical reports; SK generated the iPSCs; MB and LZ cultured and differentiated the iPSCs into neurons and characterized the cell types; MN performed the RNA sequencing analysis and PPI network analysis. MM performed the gene ontology and Hi-C analyses. DP, JF and MB researched the literature and prepared the manuscript. MB, MN, MM, SG, SuS, JF and DP edited and reviewed the manuscript, and DP and JF coordinated the writing of the manuscript. All the authors discussed, read, and approved the manuscript.

## Notes

### Competing Interest Statement

The authors have declared no competing interest.

## References

Auwerx C, Kutalik Z, Reymond A. 2024a. The pleiotropic spectrum of proximal 16p11.2 CNVs. American journal of human genetics 111: 2309–2346.

Auwerx C, Moix S, Kutalik Z, Reymond A. 2024b. Disentangling mechanisms behind the pleiotropic effects of proximal 16p11.2 BP4-5 CNVs. American journal of human genetics 111: 2347–2361.

Bhagwat SV, McMillen WT, Cai S, Zhao B, Whitesell M, Shen W, Kindler L, Flack RS, Wu W, Anderson B et al. 2020. ERK Inhibitor LY3214996 Targets ERK Pathway-Driven Cancers: A Therapeutic Approach Toward Precision Medicine. Mol Cancer Ther 19: 325–336.

Blizinsky KD, Diaz-Castro B, Forrest MP, Schurmann B, Bach AP, Martin-de-Saavedra MD, Wang L, Csernansky JG, Duan J, Penzes P. 2016. Reversal of dendritic phenotypes in 16p11.2 microduplication mouse model neurons by pharmacological targeting of a network hub. Proceedings of the National Academy of Sciences of the United States of America 113: 8520–8525.

Boucher J, Kleinridders A, Kahn CR. 2014. Insulin receptor signaling in normal and insulin-resistant states. Cold Spring Harb Perspect Biol 6.

Chang H, Li L, Li M, Xiao X. 2017. Rare and common variants at 16p11.2 are associated with schizophrenia. Schizophr Res 184: 105–108.

Chen J, Bardes EE, Aronow BJ, Jegga AG. 2009. ToppGene Suite for gene list enrichment analysis and candidate gene prioritization. Nucleic Acids Res 37: W305–311.

Chen WJ, Lin Y, Xiong ZQ, Wei W, Ni W, Tan GH, Guo SL, He J, Chen YF, Zhang QJ et al. 2011. Exome sequencing identifies truncating mutations in PRRT2 that cause paroxysmal kinesigenic dyskinesia. Nature genetics 43: 1252–1255.

Chilinski M, Sengupta K, Plewczynski D. 2022. From DNA human sequence to the chromatin higher order organisation and its biological meaning: Using biomolecular interaction networks to understand the influence of structural variation on spatial genome organisation and its functional effect. Semin Cell Dev Biol 121: 171–185.

Cremer T, Cremer M. 2010. Chromosome territories. Cold Spring Harb Perspect Biol 2: a003889.

D’Angelo D, Lebon S, Chen Q, Martin-Brevet S, Snyder LG, Hippolyte L, Hanson E, Maillard AM, Faucett WA, Mace A et al. 2016. Defining the Effect of the 16p11.2 Duplication on Cognition, Behavior, and Medical Comorbidities. JAMA psychiatry 73: 20–30.

Decker B, Liput M, Abdellatif H, Yergeau D, Bae Y, Jornet JM, Stachowiak EK, Stachowiak MK. 2020. Global Genome Conformational Programming during Neuronal Development Is Associated with CTCF and Nuclear FGFR1-The Genome Archipelago Model. Int J Mol Sci 22.

Deng W, Rupon JW, Krivega I, Breda L, Motta I, Jahn KS, Reik A, Gregory PD, Rivella S, Dean A et al. 2014. Reactivation of developmentally silenced globin genes by forced chromatin looping. Cell 158: 849–860.

Deshpande A, Yadav S, Dao DQ, Wu ZY, Hokanson KC, Cahill MK, Wiita AP, Jan YN, Ullian EM, Weiss LA. 2017. Cellular Phenotypes in Human iPSC-Derived Neurons from a Genetic Model of Autism Spectrum Disorder. Cell Rep 21: 2678–2687.

Diaz-Flores E, Goldschmidt H, Depeille P, Ng V, Akutagawa J, Krisman K, Crone M, Burgess MR, Williams O, Houseman B et al. 2013. PLC-gamma and PI3K link cytokines to ERK activation in hematopoietic cells with normal and oncogenic Kras. Sci Signal 6: ra105.

Dixon JR, Selvaraj S, Yue F, Kim A, Li Y, Shen Y, Hu M, Liu JS, Ren B. 2012. Topological domains in mammalian genomes identified by analysis of chromatin interactions. Nature 485: 376–380.

Elsayed LEO, AlHarbi NA, Alqarni AM, Eltayeb HHE, Mostafa NMM, Abdulrahim MM, Zaid HIB, Alanzi LM, Ababtain SA, Aldulaijan K et al. 2024. Chromosome 16p11.2 microdeletion syndrome with microcephaly and Dandy-Walker malformation spectrum: expanding the known phenotype. Hum Genomics 18: 95.

Finlay D, Vuori K. 2007. Novel noncatalytic role for caspase-8 in promoting SRC-mediated adhesion and Erk signaling in neuroblastoma cells. Cancer Res 67: 11704–11711.

Franke M, Daly AF, Palmeira L, Tirosh A, Stigliano A, Trifan E, Faucz FR, Abboud D, Petrossians P, Tena JJ et al. 2022. Duplications disrupt chromatin architecture and rewire GPR101-enhancer communication in X-linked acrogigantism. American journal of human genetics 109: 553–570.

Girdhar K, Hoffman GE, Bendl J, Rahman S, Dong P, Liao W, Hauberg ME, Sloofman L, Brown L, Devillers O et al. 2022. Chromatin domain alterations linked to 3D genome organization in a large cohort of schizophrenia and bipolar disorder brains. Nature neuroscience 25: 474–483.

Golzio C, Willer J, Talkowski ME, Oh EC, Taniguchi Y, Jacquemont S, Reymond A, Sun M, Sawa A, Gusella JF et al. 2012. KCTD13 is a major driver of mirrored neuroanatomical phenotypes of the 16p11.2 copy number variant. Nature 485: 363–367.

Heron SE, Dibbens LM. 2013. Role of PRRT2 in common paroxysmal neurological disorders: a gene with remarkable pleiotropy. J Med Genet 50: 133–139.

Hudac CM, Bove J, Barber S, Duyzend M, Wallace A, Martin CL, Ledbetter DH, Hanson E, Goin-Kochel RP, Green-Snyder L et al. 2020. Evaluating heterogeneity in ASD symptomatology, cognitive ability, and adaptive functioning among 16p11.2 CNV carriers. Autism Res 13: 1300–1310.

Iroegbu JD, Ijomone OK, Femi-Akinlosotu OM, Ijomone OM. 2021. ERK/MAPK signalling in the developing brain: Perturbations and consequences. Neurosci Biobehav Rev 131: 792–805.

Jacquemont S, Reymond A, Zufferey F, Harewood L, Walters RG, Kutalik Z, Martinet D, Shen Y, Valsesia A, Beckmann ND et al. 2011. Mirror extreme BMI phenotypes associated with gene dosage at the chromosome 16p11.2 locus. Nature 478: 97–102.

Jamshidi S, Rostami A, Shojaei S, Taherkhani A, Taherkhani H. 2024. Exploring natural anthraquinones as potential MMP2 inhibitors: A computational study. Biosystems 235: 105103.

Jutla A, Turner JB, Green Snyder L, Chung WK, Veenstra-VanderWeele J. 2020. Psychotic symptoms in 16p11.2 copy-number variant carriers. Autism Res 13: 187–198.

Kishi Y, Gotoh Y. 2018. Regulation of Chromatin Structure During Neural Development. Front Neurosci 12: 874.

Kumar RA, KaraMohamed S, Sudi J, Conrad DF, Brune C, Badner JA, Gilliam TC, Nowak NJ, Cook EH, Jr., Dobyns WB et al. 2008. Recurrent 16p11.2 microdeletions in autism. Human molecular genetics 17: 628–638.

Kuzniewska B, Rejmak E, Malik AR, Jaworski J, Kaczmarek L, Kalita K. 2013. Brain-derived neurotrophic factor induces matrix metalloproteinase 9 expression in neurons via the serum response factor/c-Fos pathway. Mol Cell Biol 33: 2149–2162.

Lavoie H, Gagnon J, Therrien M. 2020. ERK signalling: a master regulator of cell behaviour, life and fate. Nature reviews Molecular cell biology 21: 607–632.

Leal G, Comprido D, Duarte CB. 2014. BDNF-induced local protein synthesis and synaptic plasticity. Neuropharmacology 76 Pt C: 639–656.

Leblond CS, Le TL, Malesys S, Cliquet F, Tabet AC, Delorme R, Rolland T, Bourgeron T. 2021. Operative list of genes associated with autism and neurodevelopmental disorders based on database review. Molecular and cellular neurosciences 113: 103623.

Lee SW, Kim HK, Naidansuren P, Ham KA, Choi HS, Ahn HY, Kim M, Kang DH, Kang SW, Joe YA. 2020. Peroxidasin is essential for endothelial cell survival and growth signaling by sulfilimine crosslink-dependent matrix assembly. Faseb J 34: 10228–10241.

Leone R, Zuglian C, Brambilla R, Morella I. 2024. Understanding copy number variations through their genes: a molecular view on 16p11.2 deletion and duplication syndromes. Front Pharmacol 15: 1407865.

Lieberman-Aiden E, van Berkum NL, Williams L, Imakaev M, Ragoczy T, Telling A, Amit I, Lajoie BR, Sabo PJ, Dorschner MO et al. 2009. Comprehensive mapping of long-range interactions reveals folding principles of the human genome. Science 326: 289–293.

Liu F, Liang C, Li Z, Zhao S, Yuan H, Yao R, Qin Z, Shangguan S, Zhang S, Zou LP et al. 2023. Haplotype-specific MAPK3 expression in 16p11.2 deletion contributes to variable neurodevelopment. Brain: a journal of neurology 146: 3347–3363.

Lu L, Liu X, Huang WK, Giusti-Rodriguez P, Cui J, Zhang S, Xu W, Wen Z, Ma S, Rosen JD et al. 2020. Robust Hi-C Maps of Enhancer-Promoter Interactions Reveal the Function of Non-coding Genome in Neural Development and Diseases. Molecular cell 79: 521–534 e515.

Martino S, Carollo PS, Barra V. 2023. A Glimpse into Chromatin Organization and Nuclear Lamina Contribution in Neuronal Differentiation. Genes (Basel) 14.

McRae AM, Duncan J, Drackley A, Ing A, Allegretti V, Raski CR, Mercier A, Prada CE, Jurgensmeyer S. 2025. Further Delineation of the Proximal 16p11.2 Microdeletion Syndrome: Novel Findings Among 22 New Individuals. American journal of medical genetics Part A 197: e63873.

Medfai H, Khalil A, Rousseau A, Nuyens V, Paumann-Page M, Sevcnikar B, Furtmuller PG, Obinger C, Moguilevsky N, Peulen O et al. 2019. Human peroxidasin 1 promotes angiogenesis through ERK1/2, Akt, and FAK pathways. Cardiovasc Res 115: 463–475.

Nora EP, Lajoie BR, Schulz EG, Giorgetti L, Okamoto I, Servant N, Piolot T, van Berkum NL, Meisig J, Sedat J et al. 2012. Spatial partitioning of the regulatory landscape of the X-inactivation centre. Nature 485: 381–385.

Papantonis A, Cook PR. 2013. Transcription factories: genome organization and gene regulation. Chem Rev 113: 8683–8705.

Parnell E, Culotta L, Forrest MP, Jalloul HA, Eckman BL, Loizzo DD, Horan KKE, Dos Santos M, Piguel NH, Tai DJC et al. 2023. Excitatory Dysfunction Drives Network and Calcium Handling Deficits in 16p11.2 Duplication Schizophrenia Induced Pluripotent Stem Cell-Derived Neurons. Biological psychiatry 94: 153–163.

Pos O, Radvanszky J, Buglyo G, Pos Z, Rusnakova D, Nagy B, Szemes T. 2021. DNA copy number variation: Main characteristics, evolutionary significance, and pathological aspects. Biomed J 44: 548–559.

Qureshi AY, Mueller S, Snyder AZ, Mukherjee P, Berman JI, Roberts TP, Nagarajan SS, Spiro JE, Chung WK, Sherr EH et al. 2014. Opposing brain differences in 16p11.2 deletion and duplication carriers. J Neurosci 34: 11199–11211.

Rabiee A, Kruger M, Ardenkjaer-Larsen J, Kahn CR, Emanuelli B. 2018. Distinct signalling properties of insulin receptor substrate (IRS)-1 and IRS-2 in mediating insulin/IGF-1 action. Cell Signal 47: 1–15.

Ramosaj M, Madsen S, Maillard V, Scandella V, Sudria-Lopez D, Yuizumi N, Telley L, Knobloch M. 2021. Lipid droplet availability affects neural stem/progenitor cell metabolism and proliferation. Nature communications 12: 7362.

Rao SS, Huntley MH, Durand NC, Stamenova EK, Bochkov ID, Robinson JT, Sanborn AL, Machol I, Omer AD, Lander ES et al. 2014. A 3D map of the human genome at kilobase resolution reveals principles of chromatin looping. Cell 159: 1665–1680.

Rein B, Yan Z. 2020. 16p11.2 Copy Number Variations and Neurodevelopmental Disorders. Trends in neurosciences 43: 886–901.

Reinhard SM, Razak K, Ethell IM. 2015. A delicate balance: role of MMP-9 in brain development and pathophysiology of neurodevelopmental disorders. Frontiers in cellular neuroscience 9: 280.

Richter M, Murtaza N, Scharrenberg R, White SH, Johanns O, Walker S, Yuen RKC, Schwanke B, Bedurftig B, Henis M et al. 2019. Altered TAOK2 activity causes autism-related neurodevelopmental and cognitive abnormalities through RhoA signaling. Molecular psychiatry 24: 1329–1350.

Roskoski R, Jr. 2012. ERK1/2 MAP kinases: structure, function, and regulation. Pharmacol Res 66: 105–143.

Rovina D, La Vecchia M, Cortesi A, Fontana L, Pesant M, Maitz S, Tabano S, Bodega B, Miozzo M, Sirchia SM. 2020. Profound alterations of the chromatin architecture at chromosome 11p15.5 in cells from Beckwith-Wiedemann and Silver-Russell syndromes patients. Scientific reports 10: 8275.

Shaikh TH. 2017. Copy Number Variation Disorders. Curr Genet Med Rep 5: 183–190.

Shannon P, Markiel A, Ozier O, Baliga NS, Wang JT, Ramage D, Amin N, Schwikowski B, Ideker T. 2003. Cytoscape: a software environment for integrated models of biomolecular interaction networks. Genome Res 13: 2498–2504.

Smedley D, Haider S, Durinck S, Pandini L, Provero P, Allen J, Arnaiz O, Awedh MH, Baldock R, Barbiera G et al. 2015. The BioMart community portal: an innovative alternative to large, centralized data repositories. Nucleic Acids Res 43: W589–598.

Smith EM, Lajoie BR, Jain G, Dekker J. 2016. Invariant TAD Boundaries Constrain Cell-Type-Specific Looping Interactions between Promoters and Distal Elements around the CFTR Locus. American journal of human genetics 98: 185–201.

Sun JH, Zhou L, Emerson DJ, Phyo SA, Titus KR, Gong W, Gilgenast TG, Beagan JA, Davidson BL, Tassone F et al. 2018. Disease-Associated Short Tandem Repeats Co-localize with Chromatin Domain Boundaries. Cell 175: 224–238 e215.

Szklarczyk D, Gable AL, Lyon D, Junge A, Wyder S, Huerta-Cepas J, Simonovic M, Doncheva NT, Morris JH, Bork P et al. 2019. STRING v11: protein-protein association networks with increased coverage, supporting functional discovery in genome-wide experimental datasets. Nucleic Acids Res 47: D607–D613.

Szklarczyk D, Kirsch R, Koutrouli M, Nastou K, Mehryary F, Hachilif R, Gable AL, Fang T, Doncheva NT, Pyysalo S et al. 2023. The STRING database in 2023: protein-protein association networks and functional enrichment analyses for any sequenced genome of interest. Nucleic Acids Res 51: D638–D646.

Vangipuram M, Ting D, Kim S, Diaz R, Schule B. 2013. Skin punch biopsy explant culture for derivation of primary human fibroblasts. J Vis Exp doi:10.3791/3779: e3779.

Wendt KS, Yoshida K, Itoh T, Bando M, Koch B, Schirghuber E, Tsutsumi S, Nagae G, Ishihara K, Mishiro T et al. 2008. Cohesin mediates transcriptional insulation by CCCTC-binding factor. Nature 451: 796–801.

Won H, de la Torre-Ubieta L, Stein JL, Parikshak NN, Huang J, Opland CK, Gandal MJ, Sutton GJ, Hormozdiari F, Lu D et al. 2016. Chromosome conformation elucidates regulatory relationships in developing human brain. Nature 538: 523–527.

Yang K, Sheikh AM, Malik M, Wen G, Zou H, Brown WT, Li X. 2011. Upregulation of Ras/Raf/ERK1/2 signaling and ERK5 in the brain of autistic subjects. Genes Brain Behav 10: 834–843.

Yao B, Christian KM, He C, Jin P, Ming GL, Song H. 2016. Epigenetic mechanisms in neurogenesis. Nature reviews Neuroscience 17: 537–549.

Yoon KJ, Vissers C, Ming GL, Song H. 2018. Epigenetics and epitranscriptomics in temporal patterning of cortical neural progenitor competence. J Cell Biol 217: 1901–1914.

Zhou W, Shi Y, Li F, Wu X, Huai C, Shen L, Yi Z, He L, Liu C, Qin S. 2018. Study of the association between Schizophrenia and microduplication at the 16p11.2 locus in the Han Chinese population. Psychiatry Res 265: 198–199.

Zou H, Yu Y, Sheikh AM, Malik M, Yang K, Wen G, Chadman KK, Brown WT, Li X. 2011. Association of upregulated Ras/Raf/ERK1/2 signaling with autism. Genes Brain Behav 10: 615–624.

