## Supplemental_Fig_S1 for "16p11.2 Copy Number Variation Alters Genome Architecture and Transcriptional Regulation During Neurodevelopment"

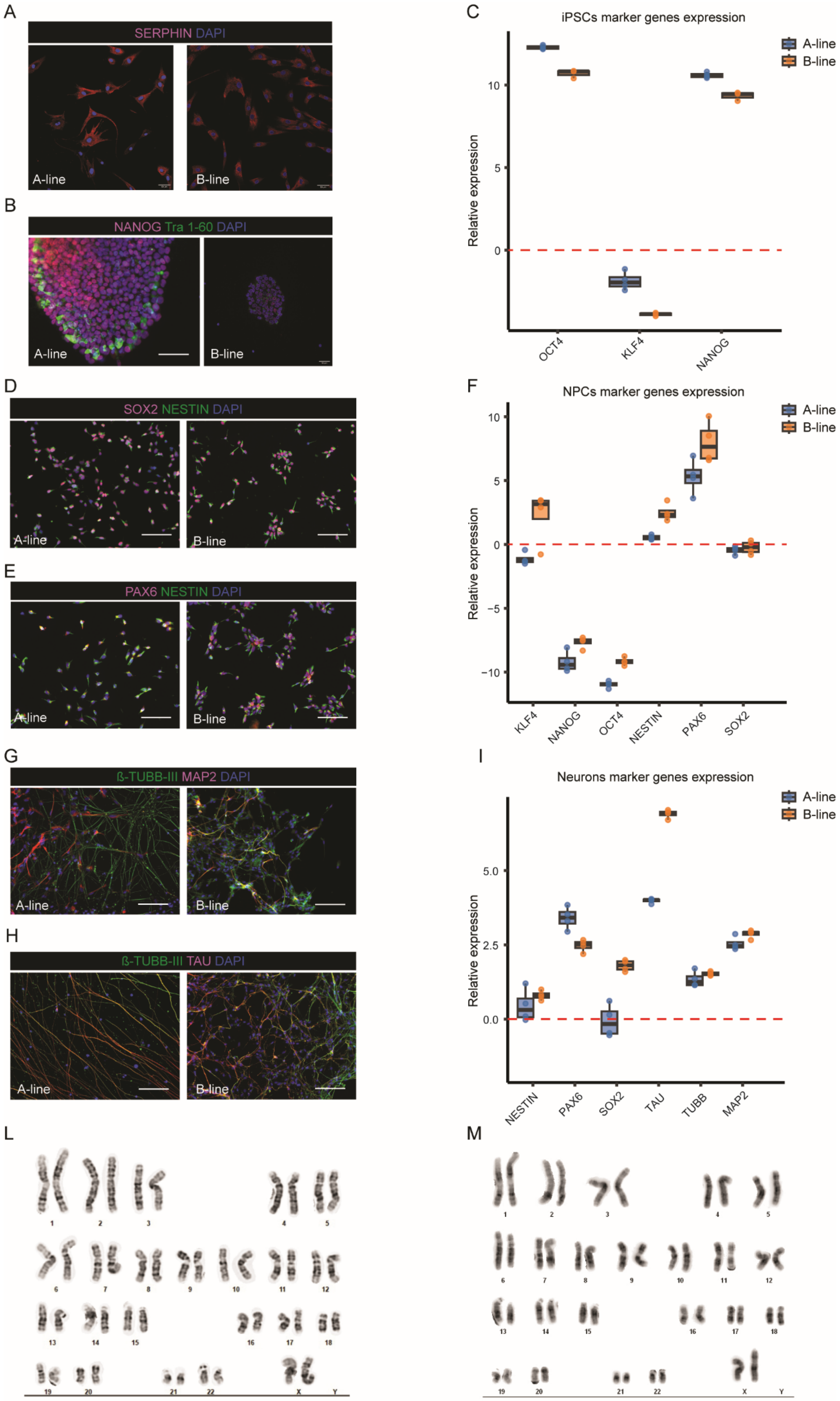
A
SERPHIN DAPI
A-line
B-line
B
NANOG Tra 1-60 DAPI
A-line
B-line
C
iPSCs marker genes expression
A-line
B-line
Relative expression
10
5
0
OCT4
KLF4
NANOG
D
SOX2 NESTIN DAPI
A-line
B-line
E
PAX6 NESTIN DAPI
A-line
B-line
F
NPCs marker genes expression
A-line
B-line
Relative expression
10
5
0
−5
−10
KLF4
NANOG
OCT4
NESTIN
PAX6
SOX2
G
ß-TUBB-III MAP2 DAPI
A-line
B-line
H
ß-TUBB-III TAU DAPI
A-line
B-line
I
Neurons marker genes expression
A-line
B-line
Relative expression
5.0
2.5
0.0
NESTIN
PAX6
SOX2
TAU
TUBB
MAP2
L
M

Supplemental Figure 1: Confirmation of the cell type identities of patient- and control-derived primary fibroblasts, iPSCs, NPCs, and neurons. A-F) Representative images of immunostaining for cell type-specific markers. A) Serphin expression was detected in human dermal fibroblasts. Scale bar = 50 µm. B) Coexpression of NANOG and TRA1-60 was detected in iPSCs. Scale bar = 50 µm. C-D) Colocalization of the nuclear NPC markers PAX6 and SOX2 in NESTIN-positive cells. Scale bar = 130 µm. E-F) Coexpression of Tubulin III with MAP2 and TAU in neuron-like cells. Scale bar = 130 µm. G-I) Boxplot showing the expression profile of cell type-specific markers analyzed by RT–qPCR. G) Expression profile of iPSC marker genes in iPSCs relative to those in fibroblasts. H) RT–qPCR analysis of iPSC and NPC gene expression in NPCs relative to that in iPSCs. I) Expression analysis of NPCs and neuronal marker genes in neurons relative to NPCs. L) Karyotype analysis of the iPSC A-line showing no chromosomal aberrations. M) Karyotype profile of the iPSC B-line showing no chromosomal aberrations.
