## Supplemental_Fig_S2 for "16p11.2 Copy Number Variation Alters Genome Architecture and Transcriptional Regulation During Neurodevelopment"

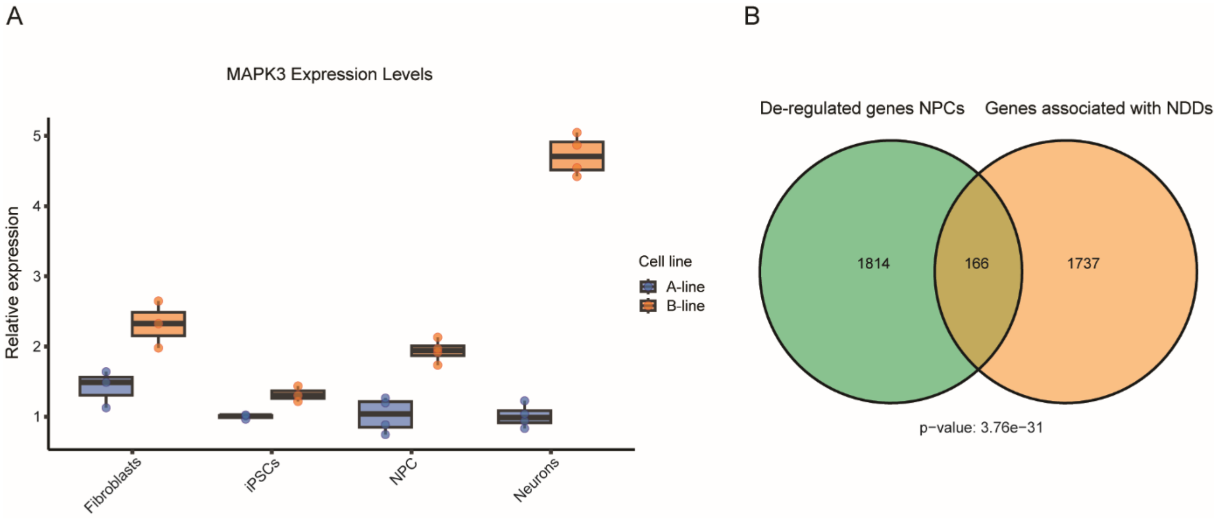


Supplemental Figure 2: A) Validation of *MAPK3* upregulation in primary fibroblasts, iPSCs, NPCs, and neurons via RT–qPCR. N=3 for fibroblasts and A-line iPSCs, and N=4 for iPSCs, NPCs, and neurons. B) Venn diagram showing the overlap of genes deregulated in patients' NPCs and genes associated with NDDs [35]. A hypergeometric test was performed; p value = 3.76e-31.
