## Supplemental_Fig_S3 for "16p11.2 Copy Number Variation Alters Genome Architecture and Transcriptional Regulation During Neurodevelopment"

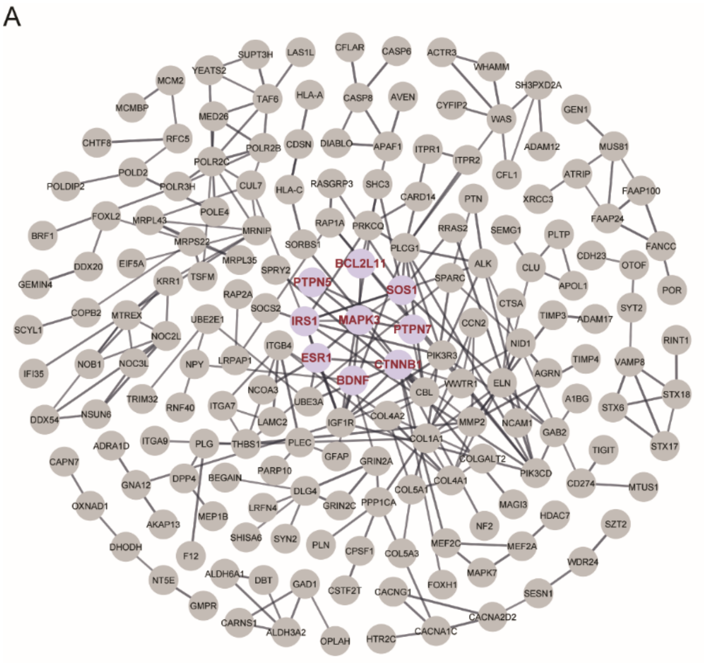


Supplemental Figure 3: Protein–protein interaction network of genes upregulated in neurons derived from the microduplication patient compared with the control.
